# Functional group-dependent responses of forest bird communities to invasive predator control and habitat fragmentation

**DOI:** 10.1101/2021.09.13.459997

**Authors:** Shaun Morgan, Nigel A. Binks, Raphael K. Didham, Andrew D. Barnes

## Abstract

**Aim:** Mounting global pressure on bird populations from invasive predators and habitat loss has driven a rapid growth in restorative and protective conservation action around the world, yet the efficacy of such actions is still not well understood. We investigated the relative effects of invasive predator control and habitat fragmentation on the abundance of native birds and invasive mammalian predators in native forest fragments.

**Location:** Waikato region, New Zealand

**Methods:** We sampled invasive mammalian predator and native bird abundances using camera traps and bird counts at 26 sites in 15 forest fragments across New Zealand’s Waikato region. Fragment area, shape complexity, and surrounding land cover of exotic and native forest were determined in ArcMap. We further created two composite gradients reflecting predator control intensity and temporal distribution of control based on seven quantitative variables recorded in each of the five years preceding native bird data collection. Finally, we estimated the relative influence of these drivers on invasive mammals and functional groups of native birds using model averaging.

**Results:** Of the two invasive predator control variables, only control intensity significantly affected invasive predator abundance and was also a more important driver than landscape or fragment structure, but responses varied among invasive mammal species. In contrast, both invasive predator control intensity and fragment structure were similarly important drivers of native bird abundance, though bird community responses varied markedly between functional groups.

**Main conclusions:** Our findings suggest that spatial extent of invasive mammal control is important for controlling mammal numbers and enhancing bird abundance, especially for small insectivorous species, and that habitat fragmentation is less important for invasive mammals but at least as important for native bird communities. Consequently, both drivers should be given strong consideration when undertaking landscape-scale conservation and restoration of bird communities in human-altered landscapes threatened by invasive predators.

## Introduction

At present, 14% of bird species are threatened with extinction and 40% are in decline due to habitat loss, predation by invasive species, overexploitation and climate change (BirdLife International, 2018; IUCN, 2019). Although habitat loss is regarded as the greatest threat to bird biodiversity (BirdLife International, 2018), invasive mammalian predators (‘invasive predators’ hereafter) are considered the greatest driver of recent extinctions, globally impacting an estimated 39% of threatened bird species (BirdLife International, 2018; Butchart, 2008).

Growing recognition of the threats posed by invasive predators to birds has driven a global expansion of conservation operations aimed at controlling or eradicating invasive predator populations on both islands and mainland regions (Jones et al., 2016). Already, studies have shown increased native bird fecundity, species richness, nesting success, survival and abundance in areas subject to invasive predator control operations, thereby demonstrating the efficacy of these methods for the conservation of native birds (Innes et al., 2010; R. K. Smith et al., 2010).

However, recent studies suggest that effective predator control may depend on the surrounding landscape context of habitat structure (Armstrong et al., 2014; García-Díaz et al., 2019; King et al., 2011). Understanding this interaction is essential for determining the regularity and intensity of predator control operations as structural landscape factors such as fragmentation and habitat connectivity influence the reinvasion potential of invasive predators such as ship rat (*Rattus rattus*) and brushtail possum (*Trichosurus vulpecula*), which can reinvade control-targeted areas from the surrounding landscape and recover to pre-control densities within just two years (Griffiths & Barron, 2016; Ji et al., 2004; King et al., 2011). Habitat structure of fragments and landscapes such as patch area, patch shape, patch isolation and land cover can alter bird abundance and community structure (Graham & Blake, 2001; Martínez-Morales, 2005). Indeed, studies have found reduced bird abundance in smaller fragments and increased nest predation across less forested landscapes (Deconchat et al., 2009; Graham & Blake, 2001). Ruffel and Didham (2017) found that benefits to native forest birds from invasive mammal control and the management of landscape forest coverage were context-dependent, yet their study did not assess the abundances of invasive mammalian predators. Additionally, an agent-based modelling study investigating the impacts of different spatiotemporal management strategies on brushtail possum populations found that control effort and spatial control distribution, rather than control timing, were the key drivers of population decline, though this study did not assess the effects on bird abundance (Lustig et al., 2019). As previous studies have typically examined these effects in isolation, an integrative approach is required to understand the relative effects of invasive predator control and landscape structure on both invasive predators and native birds, which is crucial for informing effective conservation management strategies (Ruffell & Didham, 2017).

In this study, we measure the effects of the spatiotemporal intensity of invasive predator control and the structure of both fragments and the wider landscape, hereafter ‘landscape structure’, on the abundance of native birds and invasive mammalian predators in native forest fragments. To do so, the abundances of invasive mammalian predators and native birds were sampled across 26 sites in 15 native forest fragments in New Zealand’s Waikato region. Additionally, we quantifed the variation in the intensity and temporal distribution of invasive predator control for each fragment over the five years preceding predator abundance sampling. With these data, we first tested whether invasive predator abundance decreases as the intensity and recency of predator control increases. Using a suite of functional traits collected for each sampled bird species, we quantitatively assigned species to two major functional groups and tested whether the recency and intensity of invasive predator control increased overall and functional group-specific bird abundances, bearing in mind this would also depend on the generation times and recovery rates of birds. Furthermore, we tested for the relative importance of predator control versus aspects of landscape structure (e.g. area, shape, surrounding cover) for driving the abundance of invasive predators and native birds.

## Methods

### Study area

This study was conducted in the central Waikato region of New Zealand’s North Island. The Waikato region’s temperate climate and productive soils have led to agriculture dominating regional land-use, with dairy farmland alone covering 28.7% of the region (Stats NZ, 2018). Prior to human settlement, native conifer-broadleaved forests covered an estimated 94% of the region (Ewers et al., 2006), and the country was devoid of native terrestrial mammals with the exception of three species of small bat (Atkinson, 2006). Since the settlement of the Waikato region, first by Polynesians and later by Europeans, regional native forest coverage has fallen to 21% (Landcare Research, 2015; MacLeod & Moller, 2006), and a range of invasive mammalian predators of native birds including ship rat, brushtail possum, and stoat (*Mustela erminea*) have been introduced (Atkinson, 2006). Native forest birds are particularly at risk from these invasive mammalian predators, with much evidence pointing towards mammalian predation as a major cause of species decline in native forests (Innes et al., 2010). Consequently, a range of invasive predator control operations have been established throughout the Waikato. The coverage and intensity of these operations vary greatly and, whilst fenced reserves such as Sanctuary Mountain Maungatautari receive extremely intensive ground-control, many forests like the Kaimai-Mamaku Conservation Park are effectively uncontrolled (Appendix S1).

### Site selection and experimental design

The 26 study sites were selected within 15 mixed-podocarp forest remnant fragments distributed across an 11,400 km^2^ area of the Waikato region (Fig. 1). Study sites were selected to include as wide a range as possible in levels of historical control of invasive mammalian predators, ranging from fragments that had no active control through to continuous and complete coverage of control within the past five years. Although the size of the selected patches ranged substantially from 14 hectares to over 41,000 hectares (Appendix S2), fragments were selected to be as similar as possible in three main criteria: 1) all fragments have had livestock exclusion fencing at the forest edge for at least 10 years, 2) no major tree felling occurred within the past 30 years, and 3) each fragment was at least 10 ha with a minimum width from edge to edge of 200 m.

**Figure 1.**
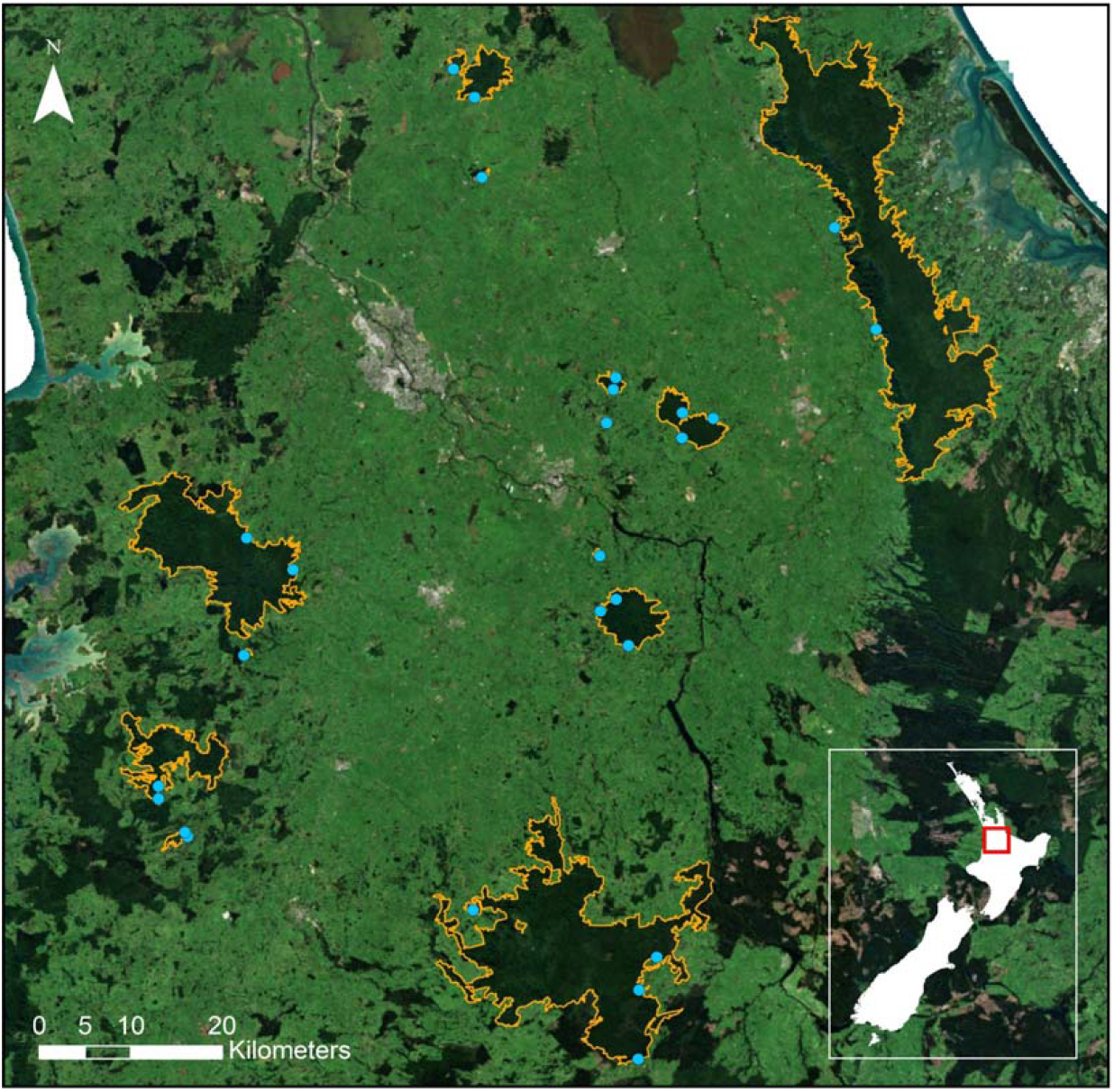
Distribution of sampled native forest fragments and individual sampling sites across the Waikato region, located in the North Island of New Zealand (see map inset). The 15 forest fragments are outlined in orange and the 26 sampling sites are represented by blue points. Image of New Zealand 10m Satellite Imagery (2017) sourced from the LINZ Data Service and licensed for reuse under the Creative Commons 4.0 New Zealand Licence.

Fragment delineation and distance/area measurements were performed using ArcMap 10.6.1 based primarily on Waikato 0.5m Rural Aerial Photos (2012-2013) and Bay of Plenty 0.4m Rural Aerial Photos (2010-2012) obtained from the LINZ data service and licensed for reuse under the Creative Commons 4.0 International Licence. Defining the fragment edges of several forested regions proved difficult as large forested areas were often tenuously connected by narrow corridors and bisected by roads. Consequently, a distinction was made between core and edge habitat to account for the potential influence of edges on habitat use by native birds and invasive mammalian predators (i.e. edge effects). Specifically, as many studies have found significantly increased nest predation along temperate forest edges (Batáry & Báldi, 2004; Vetter et al., 2013), areas of native forest fragments that narrowed to less than 200 m were considered habitat edge that lacked core habitat and were therefore treated as matrix habitat. Similarly, studies have found that narrow linear clearings created by roads can have negative impacts on the movements of understory forest birds and some small mammals (Goosem, 2001; Lees & Peres, 2009). Consequently, public roads with a lack of continuous tree canopy were treated as fragment edges.

### Invasive predator monitoring

The 26 sampling sites in which native birds and invasive predators were monitored were selected to be flat (<15° slope) and at a consistent distance of 40 m from the nearest forest edge, to spatially control for edge effects. Sampling occurred over a seven month period from December 2018 to July 2019 in an effort to avoid invasive predator control operations which are typically conducted between July and November when seasonal predator population irruptions typically occur (Elliott & Kemp, 2016).

The abundance of invasive mammalian predators was estimated using motion-sensing infra-red trail cameras (Browning Elite HD model BTC-5HDE) intermittently distributed across all 26 forest sites. Cameras were deployed at a height of 0.2 m above the ground, mounted on a wooden stake. To minimise false triggering and allow for unobstructed photographs, dense vegetation was removed where necessary to give cameras a clear line of site of at least 2 m. Each site was sampled three to five times over periods of up to 28 days, though various technical issues with the cameras meant that individual sampling periods varied from 1 to 28 days, yielding total sampling periods that varied from 38 to 117 days across the sites. Cameras were set to take five consecutive 10 MP photos over a period of 6 seconds, with a 10-second time delay between trigger events to avoid excessive numbers of events capturing the same individual that might remain at a site for an extended period. Each set of five consecutive photos was considered a single photographic event for the purposes of recording invasive predator presence/absence.

A commonly used detection threshold of 30 minutes was adopted to separate notionally ‘independent’ detection events, as past studies using this interval have typically focused on small mammals similar to the rodent and mustelid species common to New Zealand forests (Gerber et al., 2010; Nichols et al., 2017). Species counts were recorded, and three groups were selected for statistical analyses based on their high abundance and ubiquity across sites: brushtail possum, ship rat, and total mammalian predator.

### Native bird monitoring

Forest bird abundance was monitored using stationary 5-minute bird counts at each of the 40-m sampling sites. This study utilised a variation of the five-minute bird count method developed by the New Zealand Department of Scientific and Industrial Research in 1975 (Hartley, 2012). Over the 5-minute period, the number and species of birds seen or heard were recorded within a 20 m radius. During the first 3 minutes, all birds were identified and counted, whereas in the last 2 minutes, only individuals that could clearly be identified as different to those detected in the first three minutes were recorded. Point counts were conducted three times at each site over a span of three months from April to July of 2019, and only on dry days to standardise visibility and bird behaviour among sites. Only native species were retained for further analysis, and birds that could not be identified or only appeared once during the study were omitted (Appendix S1).

### Landscape structure data

The aerial imagery used to delineate study site fragments was used in conjunction with the New Zealand Land Cover Database (v4.1) in ArcMap to classify land cover within a 5 km radius of the edge of each fragment. The ‘manuka and/or kanuka’, ‘broadleaved indigenous hardwood’ and ‘indigenous forest’ classes were grouped to create a ‘native forest’ class, whilst the ‘exotic forest’ class was recorded separately. Waikato 0.5 m Rural Aerial Photos (2012-2013) and Bay of Plenty 0.4m Rural Aerial Photos (2010-2012) were again used to ground truth the Land Cover Database classification and, where necessary, polygons were reclassified.

The proportional coverage of native forest and exotic forest within 1 km and 1-5 km buffer zones around the perimeter of each fragment were then determined in ArcMap (Appendix S2). Fragment area and fractal dimension index (FDI) variables were also calculated for each of the study fragments (Appendix S2) as there is strong evidence of their influence on species abundance and dispersal success (Calabrese & Fagan, 2004; Ewers & Didham, 2007; Graham & Blake, 2001). Fragment perimeter and area geometries were calculated in ArcMap and used to determine the FDI as two times the natural logarithm of fragment perimeter (m) divided by the natural logarithm of fragment area (m^2^). FDI typically varies between 1 and 2, with higher values indicating fragment shapes with more complex perimeters (McGarigal & Marks, 1995).

### Invasive predator control data

Invasive predator control records for each study fragment and the surrounding 5 km from the edge of each fragment were obtained for a five-year period prior to the final camera monitoring day at each site. Data were collected from records obtained from landowners, land managers, and groups and organisations involved with invasive predator control operations across our study locations.

Our aim was to record a broad set of fragment-level ‘input’ variables that characterise invasive predator control efforts, from which an orthogonal set of composite variables could be created using PCA to deal with expected intercorrelation among input variables. The seven classes of input variables comprised 1) the minimum distance from each study fragment to the nearest invasive predator controlled area (including control within study fragments), along with six variables describing the proportion of each study fragment that received: 2) general invasive predator control, 3) poison-based predator control, 4) trapping-based predator control, 5) rat control, 6) possum control, and 7) both rat and possum control. Values were calculated for the 15 study fragments for each of the 5 years preceding bird monitoring, yielding a total of 35 input variables (Appendix S3).

### Constructing invasive predator control indices using Principal Component Analysis

All data analyses were performed in R version 3.6.0 (R Core Team, 2019). PCA was performed on the invasive predator control input variables (Appendix S3) with unit variance scaling using the R package ‘FactoMineR’. Based on the PCA biplot (Fig. 2), the first component was interpreted as ‘invasive predator control intensity’ due to the strong positive correlation of variables relating to the spatial coverage of predator control along the principal axis. This component explained 50.3% of the variance in the data (Figure S4). For the second component, positive vectors were generally associated with more recent predator control years (1-2), whilst negative vectors were generally associated with earlier predator control years (4-5). Consequently, the second component was interpreted as the ‘temporal distribution of invasive predator control’ and explained 17.6% of the variance (Figure S4).

**Figure 2.**
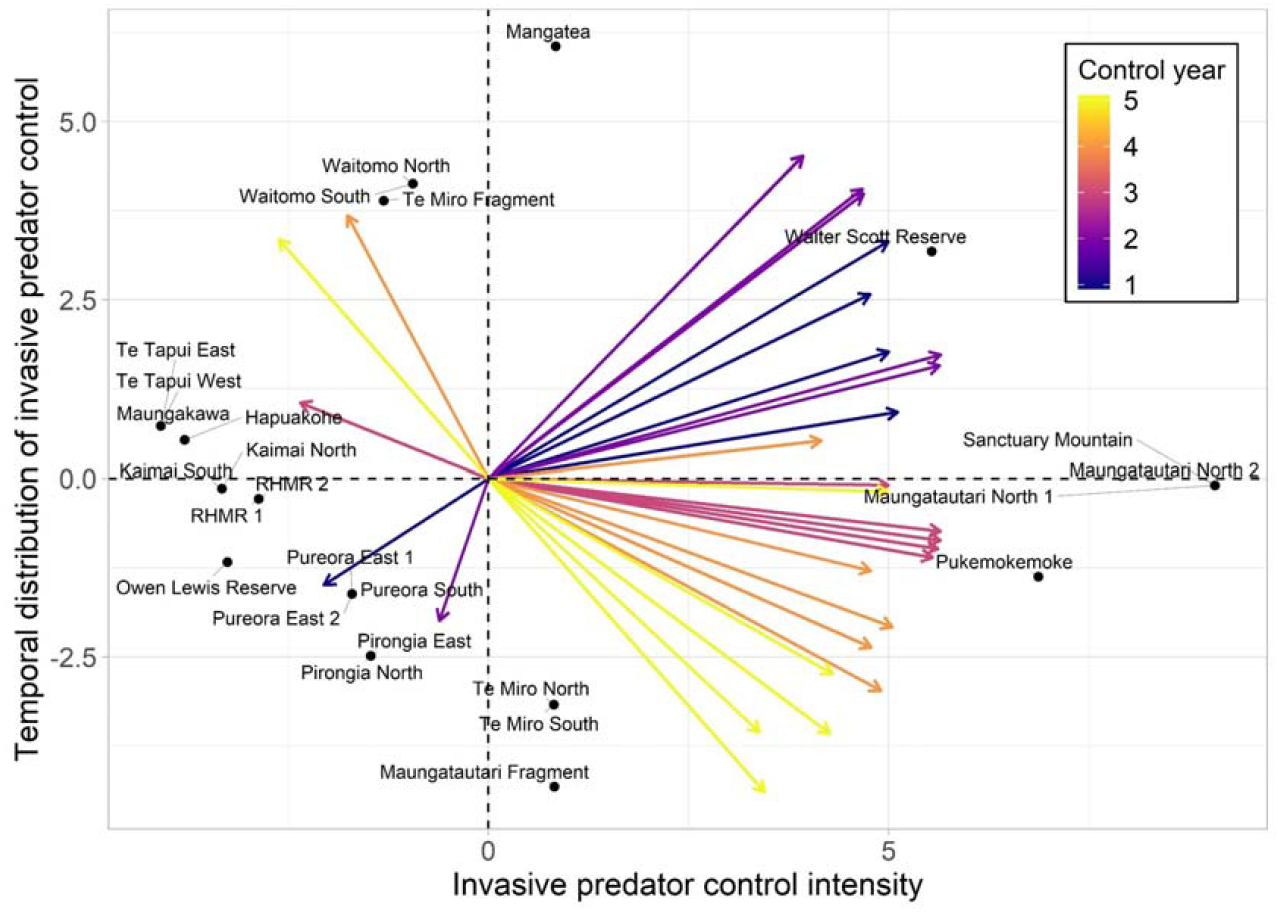
PCA biplot displaying the component scores and variable loadings obtained from the PCA of 35 variables describing invasive predator control. Labelled points represent the first (invasive predator control intensity) and second (temporal distribution of invasive predator control) PC scores for each sampling site. The direction of the vector arrows indicates the correlation of each variable with the PCs, whilst vector length indicates the strength of the contribution of each variable to the PCs. Vector colour corresponds to the number of years prior to monitoring that each invasive predator control variable is describing.

### Quantifying bird community functional groups

To examine responses in native bird communities to invasive predator control and landscape structure, bird species were first grouped by their shared functional traits (Bregman et al., 2016; Watson et al., 2004) in order to detect trait-dependent differences in species sensitivity. These functional groups were generated using the ‘dbFD’ function in the FD package (Laliberté & Legendre, 2010). To construct these groups, specific functional traits (Appendix S5) and abundances (Appendix S6) were used to create a Gower dissimilarity matrix, from which a dendrogram was created using Ward’s clustering method that revealed a distinct separation between insectivores and nectarivores (Appendix S7). The abundance of birds within these functional groups was quantified to determine the effects of invasive predator control and landscape structure on the abundances of different functional groups. Analyses were then conducted on functional groups that had sufficient numbers to detect differences across samples. Specifically, these groups included functional group 1 (hereafter, ‘nectarivores’), functional group 2 (hereafter ‘insectivores’), and the overall bird community (including all native species across the four identified functional groups; Appendix S5).

### Statistical analyses

Prior to model construction, we performed mean-centring and standard deviation-based scaling on predictor variables to enable direct comparisons of effect sizes among models. We then tested the effects of both invasive predator control indices (predator control intensity and temporal distribution of invasive predator control) on invasive predator counts (brushtail possum, ship rat and total mammalian predators) using generalised linear mixed effects models in the ‘lme4’ package. Each model contained a random effect term specifying the grouping of study sites within forest fragments, and a model offset for the (log transformed) number of monitoring days completed at each site in order to account for variation in sampling effort across sites.

To determine whether the PCA axes used to describe the intensity and temporal distribution of invasive predator control were able to explain variation in invasive mammal abundance quantified from our camera trap sampling, we tested for the effects of invasive predator control on predator counts using generalised linear regression with a negative binomial error distribution to account for overdispersion. A Likelihood Ratio Test (LRT) was undertaken using analysis of variance (ANOVA) tables to test the statistical support for more complex models containing both invasive predator control predictor variables over more simplified models based on AIC values using the ‘anova’ function in the ‘stats’ package. To test for the relative effects of invasive predator control and landscape structure on invasive mammal and native bird abundance, we constructed linear mixed effects models for each bird and invasive predator response variable using the full set of invasive predator control and landscape predictor variables. Bird response variables were modelled with a Poisson error distribution, and predator responses were modelled on a negative binomial error distribution to account for overdispersion. All models included a random effect term for the grouping of sites within fragments, and the invasive predator response models included a model offset as described above to account for variation in sampling effort across sites. Variance inflation factors (VIF) were determined for each full model and all variables were deemed independent based on a commonly used maximum threshold of 10 (O’Brien, 2007).

For each full model, the automated model selection function (‘dredge’) in the ‘MuMIn’ package was used to compute all models for every potential predictor variable combination and rank them by AIC_c_. Following Jochum et al. (2017), models with a maximum ΔAIC_c_ of 4 compared to the model with the lowest AIC_c_ were extracted for model averaging. This cut-off balanced the number of models to limit the model-averaged coefficient uncertainty associated with increasing the number of models, whilst including enough models to be fairly certain that the “best model” was not excluded (Grueber et al., 2011). The model averaging function (‘model.avg’) in the ‘MuMIn’ package was used to create the reduced model set and perform the maximum likelihood model averaging. Following the zero method (full average) of model averaging, zero value estimates were assigned to parameters that were absent from subset models before the overall averaged parameters were calculated (Burnham & Anderson, 2002). This approach decreases the effect sizes (and errors) of predictors restricted to low weighted models and is recommended when aiming to determine which factors have the strongest effect on the response variable (Nakagawa & Freckleton, 2011). The model averaging function also returned the sums of model Akaike weights for each term over all models containing that term. As in Jochum (2017), these values can be interpreted as indicators of variable importance. Akaike weighted likelihood-ratio based pseudo-R^2^ values were then calculated for each of the subset models within each averaged model set (Cox & Snell, 1989). These values were averaged to determine an R^2^ value for each of the averaged models as an indicator of model fit, and confidence intervals (95%) were determined for model parameters.

## Results

### Effects of predator control on invasive predators

Approximately 1840 days of camera-trap footage were recorded across 26 sites located in 15 separate native forest fragments across New Zealand’s Waikato region. From this footage, 957 sightings of invasive mammalian predators were recorded from seven different species, with brushtail possums and ship rats accounting for 90.1% of these observations. Model selection retained the control intensity variable in total invasive predator and brushtail possum models, but not in the ship rat model, for which the null model was selected. The temporal distribution of control variable was removed from all models.

Invasive predator control appeared to have a consistently negative effect on the observed abundance of all invasive predator classes (Fig. 3a), suggesting that, as hypothesised, increasing the intensity of control within a five-year period reduces the overall abundance of invasive mammalian predators. There was a significant negative effect of control intensity on both total invasive predator abundance (β = −0.935, z = −4.545, df = 22, P < .001) and brushtail possum abundance β = −1.013, z = −3.021, df = 22, P = .003) in single predictor models. In contrast, ship rat abundance responded more weakly to invasive predator control (Fig. 3a; β = −0.672, z = −1.927, df = 22, P = .054).

**Figure 3.**
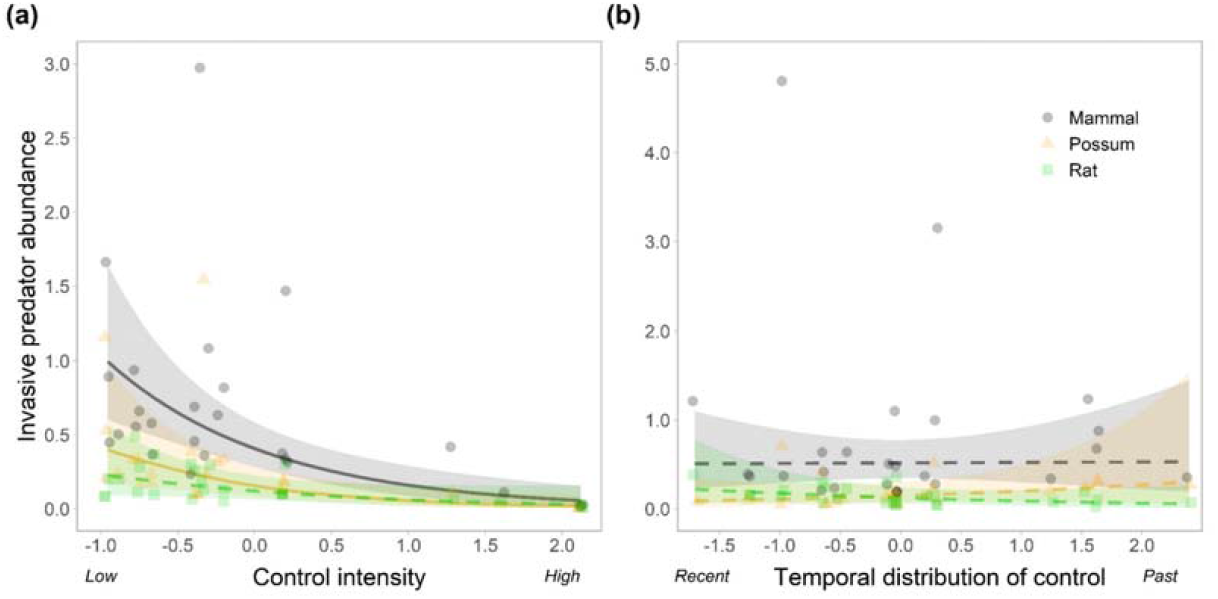
Responses of invasive predator abundances to a) the intensity of invasive predator control, and b) the temporal distribution of invasive predator control. Plots show partial effects and partial residuals for invasive predator abundance while holding other predictors at their means. Points, fitted lines, and 95% confidence intervals are grouped by abundance of all invasive predators, possums only, and rats only. Dashed lines signify a non-significant relationship (P > 0.05).

The temporal distribution of invasive predator control appeared to have only a very weak, positive effect on the total abundance of invasive predators (Fig. 3b), supporting its removal from the total invasive predator abundance model. The temporal distribution of control appeared to have weak conflicting effects on ship rat and brushtail possum abundances (Fig. 3b), and it did not have a statistically significant effect on either possum abundance (β = 0.302, z = 0.991, df = 22, P = .322) or ship rat abundance β = −0.337, z = - 1.157, df = 22, P = .247). See Appendix S8 for all model statistics.

### Relative effects of invasive predator control and landscape structure

Model averaging selected 9–51 top models for each native bird and invasive predator response variable from the full set of predictor variables describing invasive predator control and landscape structure (see Table 1 for all standardised coefficients, variable importance factors, and model inclusion counts from averaged models). Every predictor appeared at least once in every model set and R-squared values for averaged models ranged from 0.21–0.79.

**Table 1.**
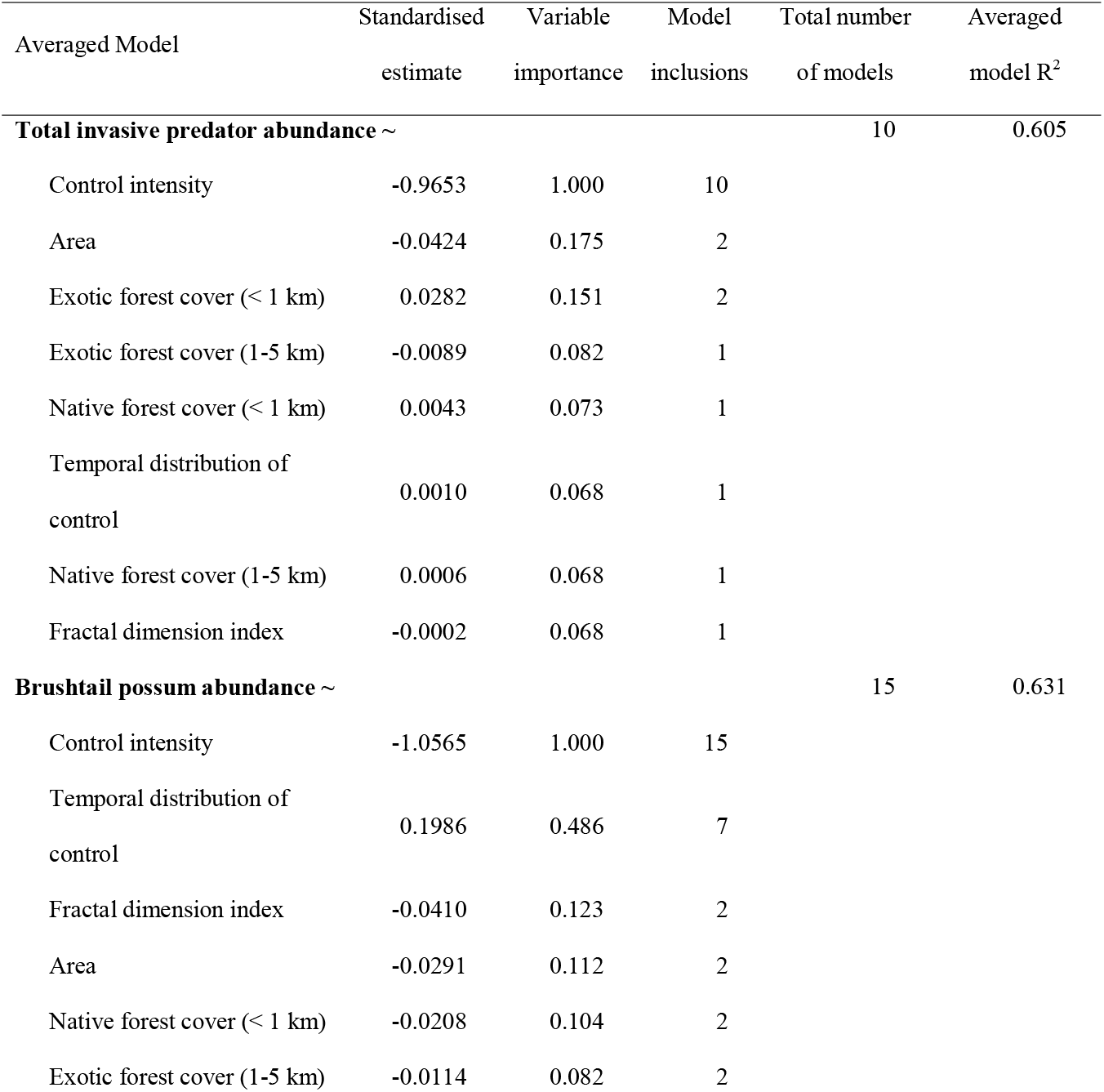
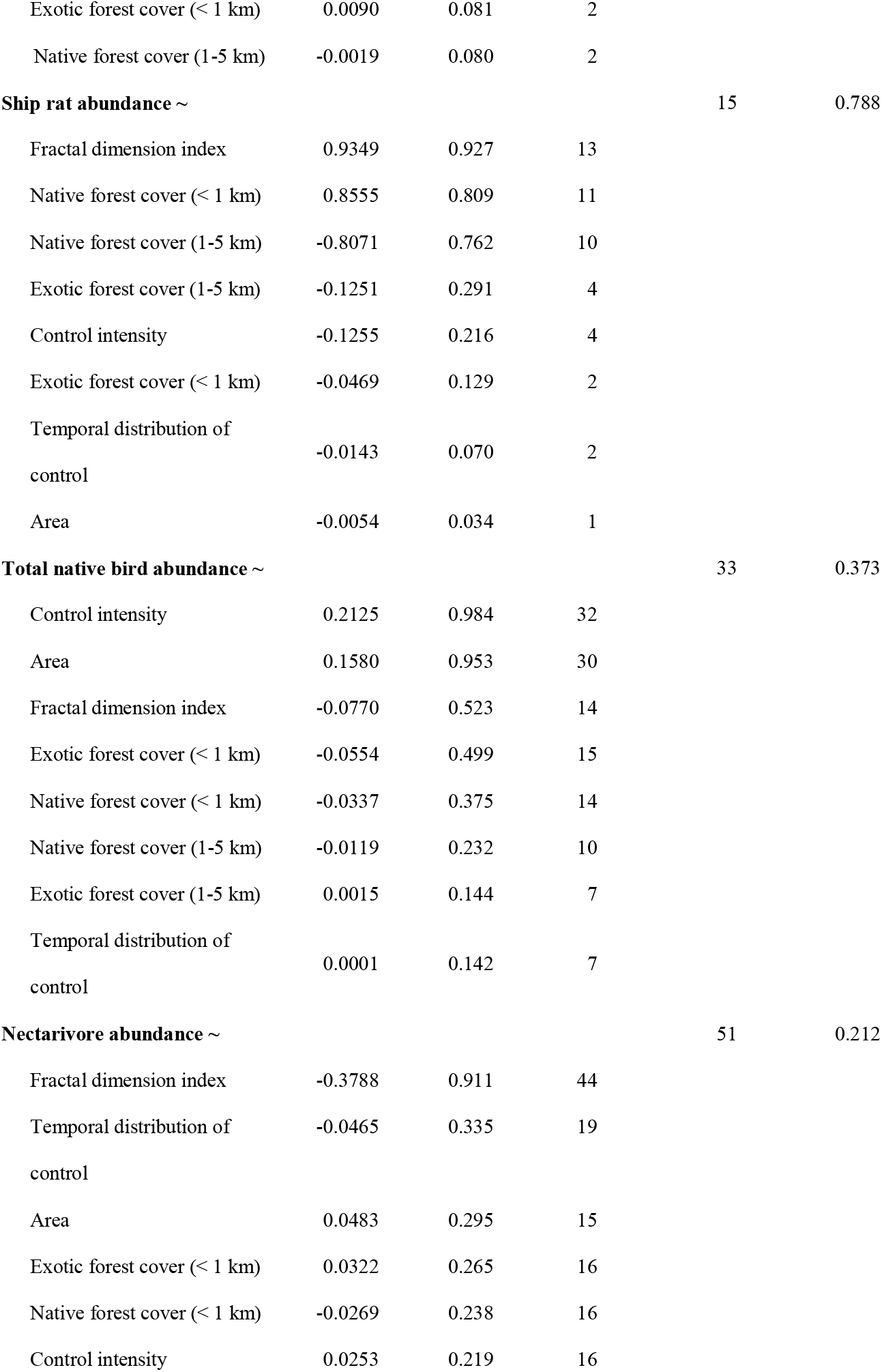
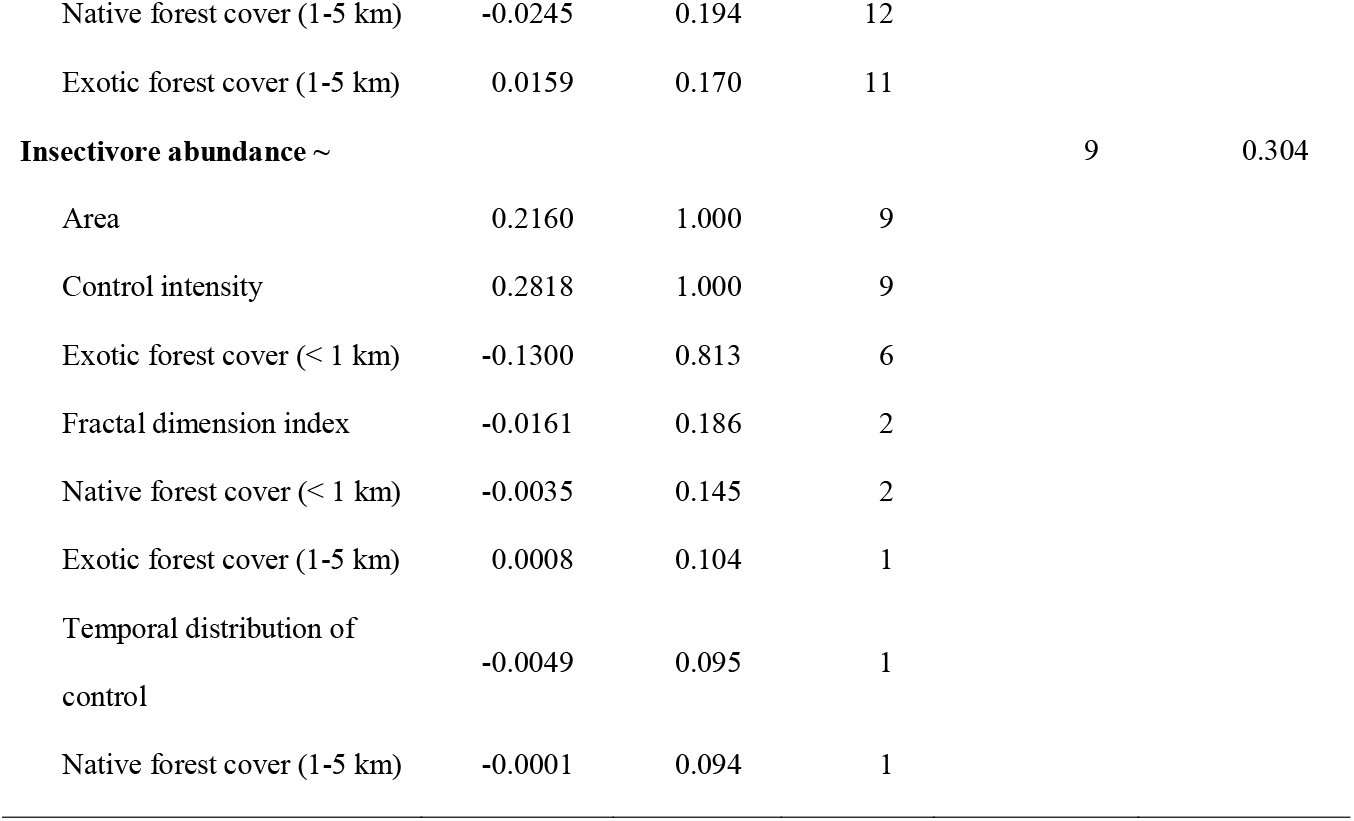
Model averaging results for invasive predator and native bird abundances. The table is split into models headed by each model’s response variable (bold). Predictor variables included in the top model set (i.e. models within ΔAICc 4 from the model with the lowest AICc) are listed beneath each response heading. Model estimates are averages of regression coefficients from the top model set, whilst variable importance gives the Akaike weight sums over all models in which the explanatory variable appears. ‘Model inclusions’ indicates the number of times each variable appears in the top model set. The total number of models in each set and the AIC-weighted average R^2^ value for each averaged model are also given.

Control intensity had a negative effect on the total abundance of invasive predators (standardised effect β = −0.965) that was greater than the effects of all landscape structure variables. Indeed, control intensity was included in all of the top models and had an effect size more than 22 times greater than the next predictor. Control intensity had a similarly strong negative effect on brushtail possum abundance (β = −1.057), though there was also a relatively consistent, albeit weak, positive effect of the temporal distribution of control (β = 0.199), which appeared in 7 of the 15 top models.

In contrast, the intensity of invasive predator control and the temporal distribution of control had relatively weak negative effects on ship rat abundance β = −0.126 and −0.014, respectively). Three landscape structure variables had similarly strong dominant effects. Both fractal dimension index and native forest cover (< 1 km) had strong positive effects on ship rat abundance (β = 0.935; 0.856), whilst a strong negative effect was associated with native forest cover (1-5 km) β = −0.807).

As expected, the effects of invasive predator control intensity on the total abundance of native birds β = 0.213) outweighed the effects of all other variables, though there was a similarly strong positive effect of fragment area β = 0.158), which was included in 30 of the 33 top models. The three next most important landscape effects were included in nearly half of the top models and had a combined effect size of similar magnitude to fragment area β = - 0.166). Interestingly, there was a strong negative effect of fractal dimension index on the abundance of nectivorous birds β = −0.379). Insectivorous bird abundances, on the other hand, responded most strongly and positively to fragment area β = 0.216) and control intensity β = 0.282), both of which appeared in every top model. Exotic forest cover (< 1 km) also had a relatively strong negative effect on insectivore abundance β = −0.130), although its effect size was less than half that of control intensity.

Of the native bird abundance models, only the confidence intervals (95%) of the control intensity, fragment area, and fractal dimension index predictors did not include zero within their ranges (Appendix S9). Partial residual plots isolating the effects of these variables showed relatively strong positive responses in total native bird abundance and insectivore abundance to control intensity and fragment area, and weaker responses in nectarivore abundance (Fig. 4a/b). The negative bird abundance responses to fractal dimension index appeared similar, though the effect on nectarivore abundance appeared slightly stronger (Fig. 4c).

**Figure 4.**
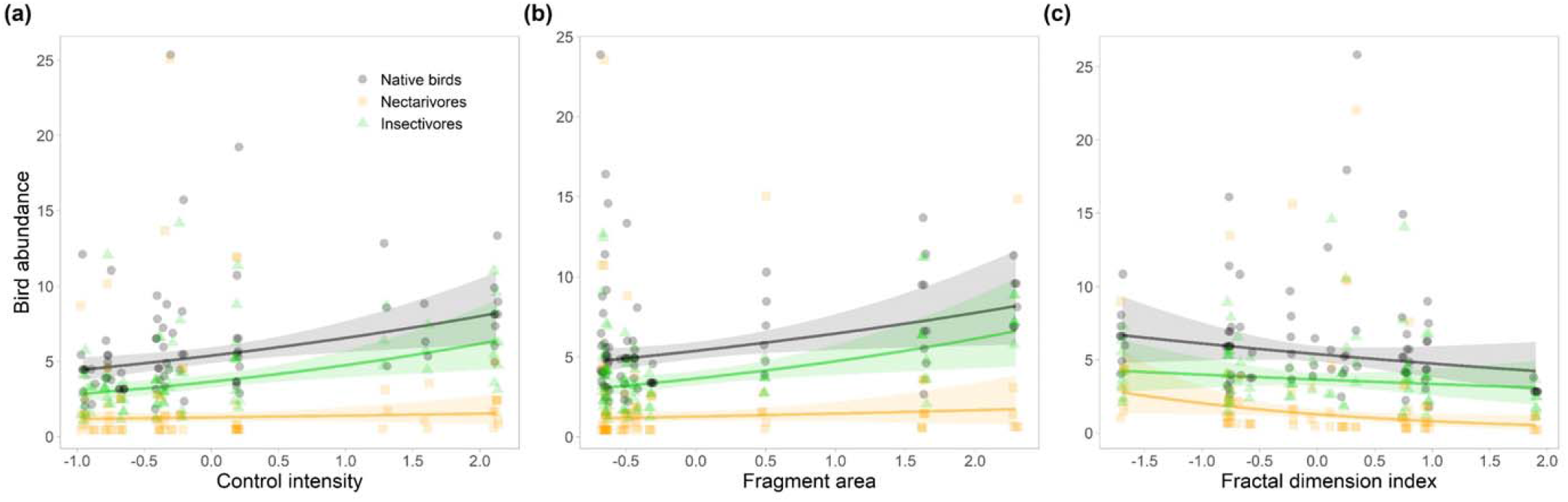
Relationships between native bird abundance classes and the most important landscape structure and predator control predictor variables in associated averaged models for which the parameter’s 95% confidence interval excluded zero. Plotted parameters include a) the intensity of invasive predator control, b) the area of study fragments, and c) the fractal dimension index of study fragments. Plots show partial effects and partial residuals for native bird abundance while holding other predictors at their means. Points, fitted lines, and 95% confidence intervals are grouped by abundance of all native birds, insectivores only, and nectarivores only.

## Discussion

We found that increased intensity of invasive predator control generally had negative effects on the abundance of invasive predators and positive effects on the abundance of native birds. Control intensity appeared to be a more important driver of invasive predator abundance than landscape structure, except with regard to ship rats, which responded much more strongly to both fragment and landscape structure. In contrast, both invasive predator control intensity and fragment structure were similarly important drivers of native bird abundance, particularly for insectivorous species. The presence of all predictors in each top model set does, however, suggest that all predictors were of some use in predicting invasive predator and native bird abundances. Overall, our findings suggest that bird conservation action in fragmented and invaded landscapes should focus simultaneously on increasing the spatial coverage of predator control and the size of forest fragments with simpler edges.

### Effects of invasive predator control on invasive predators and native birds

The dominant negative effect of invasive predator control intensity on the total abundance of invasive mammalian predators was unsurprising as many studies have demonstrated the efficacy of mammalian predator control operations (Courchamp et al., 2003; Elliott & Kemp, 2016; Van Vianen et al., 2018). However, where other studies have typically classified control intensity categorically as a function of trap/poison density and the number of active trap/poison days (Lustig et al., 2019; Ruffell et al., 2015), the continuous index used in this study was based on the proportion of fragment habitat controlled over five years. Although such scale-based indices have not been widely used, similar negative trends are still to be expected as numerous studies have suggested that increasing the size of a controlled area prolongs the effects of periodic treatment on invasive predators by reducing immigration-driven reinvasion (Brown et al., 2015; Griffiths & Barron, 2016; Lustig et al., 2019).

Interestingly, the intensity of invasive predator control appeared to strongly reduce the abundance of brushtail possums whilst enhancing the abundance of whole native bird communities and of insectivorous birds in particular. The temporal distribution of predator control also appeared to have some importance for possum abundance based on model averaging, though its effect was not statistically significant, suggesting a relatively weak influence of temporal distribution of control efforts on invasive predators in comparison to the spatial intensity of control measures.

Conversely, ship rats were not strongly affected by predator control at all. One explanation for this could be that ship rats can breed rapidly and disperse quickly across hundreds of metres of pastoral habitat to repopulate post-control fragments within months (King et al., 2011). Consequently, the use of predominantly periodic control operations across the study sites may not have been frequent enough to significantly supress populations across the monitoring period. Indeed, Ruffell et al. (2015) found that only high-intensity ongoing poison baiting could significantly reduce the abundances of rats and possums. Additionally, our models did not account for possible competitive releases of ship rat populations, which have often been observed in fragments with reduced brushtail possum abundances (Griffiths & Barron, 2016).

The strong positive effect of invasive predator control intensity on the total abundance of native birds was also expected due to the resulting reductions in predation pressure and competition for food (Innes et al., 2010). Interestingly, of the different bird functional groups, only the abundance of insectivores was significantly affected by both the intensity of invasive predator control and the total invasive predator abundance. The superior gap-crossing ability of the nectarivores (Appendix S5) could explain their lack of response to changes in predator abundance as it may enable improved predator avoidance and greater rates of patch recolonization where predation is higher (Hanski, 1998; Lees & Peres, 2009).

Our results appear to somewhat contradict previous findings that similar assemblages of insectivorous bird species were no more abundant in predator controlled sites than in uncontrolled sites, whilst nectivorous species were significantly more abundant (Innes et al., 2004; Ruffell & Didham, 2017). Ruffell and Didham (2017) proposed that the general absence of insectivore responses may have been due to ineffectual predator control or weak predation pressure. Our results did not support this assertion as invasive predator control intensity significantly reduced invasive predator abundance, and higher numbers of native birds, particularly insectivores, were found where invasive predator numbers were supressed, illustrating both the strength of predation pressure and efficacy of invasive predator control.

These contradictory findings are more likely due to differences in how predator control intensity was defined. Whilst studies such as Ruffell and Didham (2017) have classified predator control intensity in a categorical manner based primarily on poison/trapping density and application frequency (sub-annual and annual), this study utilised a continuous index based on the spatial coverage of predator control within isolated fragments. In fact, the former is more closely related to our index of the temporal distribution of control which was similarly found to have very little effect on the abundance of birds and invasive predators in our study. Taken together, our results suggest that increasing the spatial coverage of invasive predator control may be considerably more important for reducing invasive mammal numbers and enhancing native bird abundance, particularly with respect to insectivores.

The lack of significant effects of the timing of invasive predator control on any response variable was unexpected as post-control population recovery of mammalian predators and native birds is generally observable within a 0-5 year window (Armstrong, 2017; Griffiths & Barron, 2016; Le Corre et al., 2015; Van Vianen et al., 2018). This suggests that either the temporal interval was too large to capture the effects of sub-annual control on predator species capable of more rapid population recovery (e.g. ship rats), or that the effects of predator control may last more than 5 years (i.e. the time span of the study was too short) (Van Vianen et al., 2018). In other words, following an initial post-control population decline, predator populations may increase to pre-control levels within one year, or they may remain at relatively low levels for more than 5 years with native bird populations already recovered.

### Invasive predator control impacts predators more than landscape structure

As we expected based on the strong effects of invasive predator control detected in the initial linear models, the intensity of invasive predator control was a more dominant driver of total invasive predator abundance than landscape structure. Control intensity was a similarly dominant driver of brushtail possum abundance, although the temporal distribution of control was also relatively important when compared to landscape variables. Curiously, the positive temporal effect suggests that possum numbers are greater in fragments under more recent control, whilst typical population recovery lag-times suggest that, if anything, the opposite trend should exist (e.g. Ji et al., 2004). A possible explanation for this is that more recent control operations have been less effective at reducing possum abundances than more historic ones, though a more detailed investigation is needed to better understand this trend.

In contrast, ship rats were far more responsive to landscape variables than either predator control variable, which is in keeping with past studies that demonstrate sensitivity of rodents and other small mammals to structural landscape features (Crooks, 2002; Presley et al., 2019). Interestingly, the proportion of native forest within both 1 kilometre, and 1 to 5 kilometre ranges from the study fragments were highly important, yet the two variables had opposing effects on ship rat abundance, suggesting that the effect of native forest coverage is scale-dependent; a phenomenon that has been demonstrated across a range of scales for a variety of species and landscapes (Deconchat et al., 2009; A. C. Smith et al., 2011). Whilst our results imply that surrounding native habitat may be somewhat important for recolonising fragments, the negative effects of native forest cover at 1 to 5 kilometres indicate that there are complex landscape dynamics that require further targeted investigations.

Fractal dimension index also had an important positive effect on ship rat abundance, suggesting that fragments with complex edges and high edge-to-area ratios support greater ship rat populations. Generally, fragments with complex edges tend to have greater population turnover and reduced population stability due to an increase in the likelihood of an individual encountering an edge (Ewers & Didham, 2006). However, fragments with greater proportions of edge habitat are also more strongly influenced by edge effects which can lead to increases in the abundance of habitat generalist species (Ewers & Didham, 2006), thereby providing a potential explanation for this positive effect on ship rats. Our findings demonstrate the considerable variability in how invasive predator species can respond to control measures and landscape structure.

### Invasive predator control and landscape structure jointly influence native bird abundance

Along with invasive predator abundance, the two fragment-specific variables in this study, area and fractal dimension index were the most consistently important predictors of native bird abundance across each class. Indeed, mammalian predator invasion and landscape structure have both previously been shown to impact a variety of avian species (Bregman et al., 2014; Innes et al., 2010; Jones et al., 2016).

The negative effect of fractal dimension index on total bird abundance contrasts with its positive effect on ship rat abundance, suggesting that the negative response of native birds to fragments with greater proportions of edge habitat may be due to increased nest predation by ship rats. Such an effect is, however, typically contextually dependent on a broader range of landscape factors and species-specific relationships (Lahti, 2001; Vetter et al., 2013). This appeared to be the case in our study as fractal dimension index exhibited a much stronger relative effect on the abundance of nectarivores than insectivores. Our results suggest that nectivorous birds could be much more sensitive to factors relating to landscape structure than to invasive predators. The relatively low value of the averaged model’s AIC-weighted R^2^ value (R^2^ = 0.212) does, however, suggest that other important variables likely contribute to variation in the abundance of nectivorous birds.

The vulnerability of insectivores to predation and habitat loss is highlighted by the equally dominant positive effects of control intensity and fragment area on insectivore abundance. While increasing habitat area typically has a positive effect on bird communities, the minimum area required to sustain forest bird populations is highly species-dependent (Innes et al., 2010; Watson et al., 2004). Terrestrial insectivore populations tend to be particularly area-sensitive, likely due to their typically limited inter-patch dispersal abilities (Martensen et al., 2008). The invertebrate food resources of ground-foraging insectivores have also been found to decline significantly with decreasing fragment size (Zanette et al., 2000). Consequently, insectivores in smaller fragments are likely to suffer greater bottom-up control due to a more limited food supply (Karr et al., 1992). Interestingly, the landscape coverage of exotic forest (< 1 km) also had a relatively important negative effect on the abundance of insectivores. This effect is somewhat expected as lower native bird abundances have generally been recorded in New Zealand’s exotic pine forests than in native forests due to reduced provisions of food and nesting habitat (Clout & Gaze, 1984; Deconchat et al., 2009).

Overall, the intensity of invasive predator control and landscape structure jointly influence native bird abundance, though the relative importance of specific aspects of landscape structure is contextually dependent.

## Conclusions

We demonstrate the importance of both predator control and landscape structure for the conservation of native birds in native forest fragments. In particular, our results suggest that the annual spatial coverage of invasive predator control operations is more important for native bird conservation than the temporal distribution of said operations across the preceding five years. At the same time, we show that conservation efforts should also focus on preventing further reduction in the size of habitat fragments and expansion of habitat edges. That said, we show significant variation in species-specific responses of invasive predators and functional group-specific responses of native birds to predator control and landscape structure. Our results indicate that, whilst currently employed control measures can effectively suppress invasive predator populations, the structure of habitat fragments and overall landscape cover also play an important role in invasive predator numbers. Similarly varying effects of these factors were evident across the bird functional groups, with insectivores responding more strongly to fragment area and control intensity, and nectarivores responding primarily to the shape complexity of habitat fragments. Consequently, strategies for conserving and restoring native bird communities in predator-invaded fragmented landscapes need to account for taxon-dependent and functional group-dependent responses of invasive predators and native birds to invasive species control measures and landscape structure.

## Acknowledgements

For providing us with historical records of invasive predator control, our thanks goes to Andrew McConnell and the Biosecurity team at Waikato Regional Council, Raymond Tana and Julia Loepelt from the Department of Conservation, Melissa Sinton from the QEII National Trust, Daniel Howie from the Maungatautari Ecological Island Trust, Steve McClunie and Dianne June from the Mount Pirongia Restoration Society, David Peters and Emma Cronin from the Aongatete Forest Restoration Trust, Colleen Grayling from the Howick Tramping Club, Phillip Dawson from OSPRI New Zealand, and Neil Fitzgerald from Forest & Bird. This project was supported by the University of Waikato Strategic Research Fund.

## Data accessibility

The data that supports the findings of this study are available in the supplementary material of this article

