## Appendix for "Functional group-dependent responses of forest bird communities to invasive predator control and habitat fragmentation"

### Appendices

**Appendix S1.** Standardised measures of invasive predator control (PCA derived), invasive predator abundance and the number of associated camera trapping days, and native bird abundance at each study site.

| Site | Control intensity | Temporal distribution of control | Total invasive predator abundance | Brushtail possum abundance | Ship rat abundance | No. camera trapping days | Total native bird abundance | Nectarivore abundance | Insectivore abundance |
| --- | --- | --- | --- | --- | --- | --- | --- | --- | --- |
| Hapuakohe | -0.886 | 0.212 | 20 | 8 | 11 | 59 | 8 | 0 | 7 |
| Kaimai North | -0.777 | -0.054 | 32 | 21 | 8 | 67 | 18 | 1 | 17 |
| Kaimai South | -0.777 | -0.054 | 68 | 24 | 41 | 75 | 20 | 6 | 14 |
| Mangatea | 0.197 | 2.393 | 21 | 15 | 6 | 67 | 11 | 0 | 10 |
| Maungakawa | -0.956 | 0.289 | 9 | 9 | 0 | 46 | 13 | 4 | 9 |
| Maungatautari Fragment | 0.193 | -1.705 | 75 | 7 | 64 | 79 | 20 | 6 | 13 |
| Maungatautari North 1 | 2.120 | -0.037 | 0 | 0 | 0 | 52 | 22 | 4 | 18 |
| Maungatautari North 2 | 2.120 | -0.037 | 0 | 0 | 0 | 52 | 32 | 13 | 17 |
| Owen Lewis Reserve | -0.762 | -0.461 | 57 | 6 | 38 | 91 | 14 | 3 | 11 |
| Pirongia East | -0.343 | -0.981 | 29 | 6 | 12 | 84 | 21 | 11 | 9 |
| Pirongia North | -0.343 | -0.981 | 63 | 55 | 8 | 38 | 12 | 3 | 8 |
| Pukemokemoke | 1.605 | -0.542 | 7 | 2 | 5 | 63 | 16 | 4 | 12 |
| Pureora East 1 | -0.398 | -0.637 | 3 | 1 | 2 | 70 | 8 | 2 | 4 |
| Pureora East 2 | -0.398 | -0.637 | 47 | 1 | 44 | 117 | 23 | 4 | 16 |
| Pureora South | -0.398 | -0.637 | 48 | 11 | 26 | 77 | 17 | 3 | 11 |
| RHMR 1 | -0.670 | -0.111 | 13 | 8 | 4 | 69 | 6 | 2 | 4 |
| RHMR 2 | -0.670 | -0.111 | 48 | 6 | 30 | 95 | 7 | 1 | 5 |
| Sanctuary Mountain | 2.120 | -0.037 | 1 | 0 | 1 | 76 | 35 | 10 | 25 |
| Te Miro Fragment | -0.305 | 1.537 | 77 | 20 | 48 | 79 | 22 | 3 | 12 |
| Te Miro North | 0.191 | -1.251 | 19 | 13 | 5 | 56 | 12 | 1 | 10 |
| Te Miro South | 0.191 | -1.251 | 21 | 10 | 5 | 56 | 23 | 4 | 17 |
| Te Tapui East | -0.956 | 0.289 | 112 | 104 | 3 | 77 | 21 | 10 | 10 |
| Te Tapui West | -0.956 | 0.289 | 36 | 35 | 0 | 42 | 8 | 4 | 4 |
| Waitomo North | -0.220 | 1.631 | 70 | 60 | 4 | 88 | 20 | 5 | 13 |
| Waitomo South | -0.220 | 1.631 | 58 | 57 | 0 | 88 | 12 | 2 | 10 |
| Walter Scott Reserve | 1.294 | 1.256 | 23 | 13 | 6 | 77 | 17 | 5 | 12 |

**Appendix S2.** Location and structural properties of native forest fragments and associated sample sites. Habitat classes describe the percentage landscape coverage of native and exotic forest within 1 and 1-5 km of the edge of each study fragment. Landcover data were obtained from the New Zealand Land Cover Database v4.1 (Landcare Research, 2015).

| Fragment | Site | Northing | Easting | Area (ha) | Edge (km) | Fractal Dimension Index | Exotic forest % (1-5 km) | Native forest % (1-5 km) | Exotic forest % (< 1 km) | Native forest % (< 1 km) |
| --- | --- | --- | --- | --- | --- | --- | --- | --- | --- | --- |
| Hapuakohe | Hapuakohe | 5846136.52 | 1808440.67 | 1931.71 | 47.54 | 1.28 | 2.10 | 13.94 | 1.47 | 13.87 |
| Kaimai | Kaimai North | 5831569.61 | 1847872.69 | 41423.75 | 407.42 | 1.30 | 5.59 | 13.13 | 9.45 | 10.29 |
| Kaimai | Kaimai South | 5820171.01 | 1852396.18 | 41423.75 | 407.42 | 1.30 | 5.59 | 13.13 | 9.45 | 10.29 |
| Mangatea | Mangatea | 5849277.99 | 1806123.13 | 287.67 | 11.58 | 1.26 | 1.55 | 24.61 | 0.09 | 18.53 |
| Maungatautari Fragment | Maungatautari Fragment | 5794847.40 | 1822148.07 | 44.78 | 4.27 | 1.28 | 1.26 | 6.51 | 0.67 | 12.76 |
| Maungatautari Mountain | Maungatautari North 1 | 5789987.36 | 1823928.62 | 3237.25 | 38.99 | 1.22 | 2.67 | 6.05 | 0.79 | 7.77 |
| Maungatautari Mountain | Maungatautari North 2 | 5788668.29 | 1822220.95 | 3237.25 | 38.99 | 1.22 | 2.67 | 6.05 | 0.79 | 7.77 |
| Maungatautari Mountain | Sanctuary Mountain | 5784812.84 | 1825290.97 | 3237.25 | 38.99 | 1.22 | 2.67 | 6.05 | 0.79 | 7.77 |
| Owen Lewis Reserve | Owen Lewis Reserve | 5755235.28 | 1808262.29 | 149.26 | 8.97 | 1.28 | 1.42 | 43.07 | 2.54 | 25.54 |
| Pirongia | Pirongia East | 5793292.81 | 1788556.38 | 16128.84 | 162.40 | 1.27 | 6.32 | 15.10 | 11.08 | 13.69 |
| Pirongia | Pirongia North | 5796856.16 | 1783474.18 | 16128.84 | 162.40 | 1.27 | 6.32 | 15.10 | 11.08 | 13.69 |
| Pukemokemoke | Pukemokemoke | 5837167.83 | 1809244.98 | 43.23 | 3.94 | 1.28 | 1.36 | 1.31 | 12.69 | 7.77 |
| Pureora | Pureora East 1 | 5746299.39 | 1826447.04 | 32231.61 | 365.09 | 1.31 | 18.16 | 15.56 | 13.61 | 14.83 |
| Pureora | Pureora East 2 | 5749980.29 | 1828369.43 | 32231.61 | 365.09 | 1.31 | 18.16 | 15.56 | 13.61 | 14.83 |
| Pureora | Pureora South | 5738560.73 | 1826354.07 | 32231.61 | 365.09 | 1.31 | 18.16 | 15.56 | 13.61 | 14.83 |
| RHMR | RHMR 1 | 5767685.48 | 1773804.39 | 4833.08 | 138.15 | 1.34 | 4.18 | 32.33 | 6.50 | 16.24 |
| RHMR | RHMR 2 | 5769092.88 | 1773824.60 | 4833.08 | 138.15 | 1.34 | 4.18 | 32.33 | 6.50 | 16.24 |
| Te Miro | Te Miro North | 5814778.74 | 1823884.27 | 403.58 | 13.67 | 1.25 | 2.50 | 7.09 | 1.47 | 5.00 |
| Te Miro | Te Miro South | 5813446.94 | 1823624.54 | 403.58 | 13.67 | 1.25 | 2.50 | 7.09 | 1.47 | 5.00 |
| Te Miro Fragment | Te Miro Fragment | 5809698.36 | 1822877.63 | 13.98 | 2.06 | 1.29 | 1.56 | 12.47 | 0.87 | 8.00 |
| Te Tapui | Maungakawa | 5810849.97 | 1831197.04 | 2326.19 | 41.03 | 1.25 | 1.71 | 3.59 | 1.25 | 2.50 |
| Te Tapui | Te Tapui East | 5810217.05 | 1834577.16 | 2326.19 | 41.03 | 1.25 | 1.71 | 3.59 | 1.25 | 2.50 |
| Te Tapui | Te Tapui West | 5808022.72 | 1831153.70 | 2326.19 | 41.03 | 1.25 | 1.71 | 3.59 | 1.25 | 2.50 |
| Waitomo | Waitomo North | 5763952.34 | 1776708.86 | 249.72 | 14.52 | 1.30 | 9.44 | 32.21 | 0.69 | 9.38 |
| Waitomo | Waitomo South | 5763444.03 | 1777007.80 | 249.72 | 14.52 | 1.30 | 9.44 | 32.21 | 0.69 | 9.38 |
| Walter Scott Reserve | Walter Scott Reserve | 5783719.33 | 1783169.87 | 48.74 | 3.71 | 1.26 | 6.30 | 27.56 | 21.21 | 21.53 |

**Appendix S3.** Aspects of fragment-level invasive predator control used to construct principle components (PCs) describing the intensity and temporal distribution of control within each study fragment. Distances are to the nearest controlled area from each study fragment’s edge (including the fragment itself), and proportions are of each site’s fragment area. Variables describe control operations that took place across each of the 5 years prior to this study (2014-2019) (e.g. y4 = invasive predator control across the fourth year prior to the final mammal or bird abundance sample taken at each site).

| Fragment site | Distance to nearest control (km) | | | | | Total controlled proportion (%) | | | | | Poison controlled proportion (%) | | | | | Trapping controlled proportion (%) | | | | |
| --- | --- | --- | --- | --- | --- | --- | --- | --- | --- | --- | --- | --- | --- | --- | --- | --- | --- | --- | --- | --- |
|  | y1 | y2 | y3 | y4 | y5 | y1 | y2 | y3 | y4 | y5 | y1 | y2 | y3 | y4 | y5 | y1 | y2 | y3 | y4 | y5 |
| Hapuakohe | 0.17 | 0.17 | 6.70 | 6.70 | 0.01 | 0.00 | 0.00 | 0.00 | 0.00 | 0.00 | 0.00 | 0.00 | 0.00 | 0.00 | 0.00 | 0.00 | 0.00 | 0.00 | 0.00 | 0.00 |
| Kaimai North | 0.00 | 0.00 | 0.00 | 0.00 | 0.00 | 1.11 | 1.11 | 1.62 | 1.11 | 1.11 | 1.11 | 1.11 | 1.62 | 1.11 | 1.11 | 1.62 | 1.62 | 1.62 | 1.11 | 1.11 |
| Kaimai South | 0.00 | 0.00 | 0.00 | 0.00 | 0.00 | 1.11 | 1.11 | 1.62 | 1.11 | 1.11 | 1.11 | 1.11 | 1.62 | 1.11 | 1.11 | 1.62 | 1.62 | 1.62 | 1.11 | 1.11 |
| Mangatea | 0.00 | 0.00 | 9.56 | 9.56 | 3.39 | 100.00 | 100.00 | 0.00 | 0.00 | 0.00 | 100.00 | 100.00 | 0.00 | 0.00 | 0.00 | 100.00 | 100.00 | 0.00 | 0.00 | 0.00 |
| Maungakawa | 1.37 | 0.00 | 14.54 | 4.03 | 4.03 | 0.00 | 0.00 | 0.00 | 0.00 | 0.00 | 0.00 | 0.00 | 0.00 | 0.00 | 0.00 | 0.00 | 0.00 | 0.00 | 0.00 | 0.00 |
| Maungatautari Fragment | 3.99 | 3.99 | 0.00 | 0.00 | 0.00 | 0.00 | 0.00 | 100.00 | 100.00 | 100.00 | 0.00 | 0.00 | 100.00 | 100.00 | 100.00 | 0.00 | 0.00 | 0.00 | 0.00 | 0.00 |
| Maungatautari North 1 | 0.00 | 0.00 | 0.00 | 0.00 | 0.00 | 100.00 | 100.00 | 100.00 | 100.00 | 100.00 | 100.00 | 100.00 | 100.00 | 100.00 | 100.00 | 100.00 | 100.00 | 100.00 | 100.00 | 100.00 |
| Maungatautari North 2 | 0.00 | 0.00 | 0.00 | 0.00 | 0.00 | 100.00 | 100.00 | 100.00 | 100.00 | 100.00 | 100.00 | 100.00 | 100.00 | 100.00 | 100.00 | 100.00 | 100.00 | 100.00 | 100.00 | 100.00 |
| Owen Lewis Reserve | 4.33 | 4.33 | 4.33 | 0.00 | 0.11 | 0.00 | 0.00 | 0.00 | 39.04 | 0.00 | 0.00 | 0.00 | 0.00 | 39.04 | 0.00 | 0.00 | 0.00 | 0.00 | 39.04 | 0.00 |
| Pirongia East | 0.00 | 0.00 | 0.00 | 0.00 | 0.00 | 6.22 | 6.22 | 6.22 | 6.22 | 100.00 | 6.22 | 6.22 | 6.22 | 6.22 | 100.00 | 0.00 | 0.00 | 0.00 | 0.00 | 5.91 |
| Pirongia North | 0.00 | 0.00 | 0.00 | 0.00 | 0.00 | 6.22 | 6.22 | 6.22 | 6.22 | 100.00 | 6.22 | 6.22 | 6.22 | 6.22 | 100.00 | 0.00 | 0.00 | 0.00 | 0.00 | 5.91 |
| Pukemokemoke | 0.00 | 0.00 | 0.00 | 0.00 | 0.00 | 100.00 | 100.00 | 100.00 | 100.00 | 100.00 | 100.00 | 100.00 | 100.00 | 100.00 | 100.00 | 0.00 | 0.00 | 0.00 | 0.00 | 0.00 |
| Pureora East 1 | 0.00 | 0.00 | 0.00 | 0.00 | 0.00 | 18.76 | 15.36 | 29.93 | 10.29 | 95.00 | 18.76 | 15.36 | 29.93 | 10.29 | 95.00 | 0.06 | 0.06 | 0.00 | 0.06 | 0.06 |
| Pureora East 2 | 0.00 | 0.00 | 0.00 | 0.00 | 0.00 | 18.76 | 15.36 | 29.93 | 10.29 | 95.00 | 18.76 | 15.36 | 29.93 | 10.29 | 95.00 | 0.06 | 0.06 | 0.00 | 0.06 | 0.06 |
| Pureora South | 0.00 | 0.00 | 0.00 | 0.00 | 0.00 | 18.76 | 15.36 | 29.93 | 10.29 | 95.00 | 18.76 | 15.36 | 29.93 | 10.29 | 95.00 | 0.06 | 0.06 | 0.00 | 0.06 | 0.06 |
| RHMR 1 | 3.70 | 0.00 | 5.49 | 0.00 | 5.49 | 0.00 | 1.28 | 0.00 | 72.07 | 0.00 | 0.00 | 1.28 | 0.00 | 72.07 | 0.00 | 0.00 | 1.28 | 0.00 | 72.07 | 0.00 |
| RHMR 2 | 3.70 | 0.00 | 5.49 | 0.00 | 5.49 | 0.00 | 1.28 | 0.00 | 72.07 | 0.00 | 0.00 | 1.28 | 0.00 | 72.07 | 0.00 | 0.00 | 1.28 | 0.00 | 72.07 | 0.00 |
| Sanctuary Mountain | 0.00 | 0.00 | 0.00 | 0.00 | 0.00 | 100.00 | 100.00 | 100.00 | 100.00 | 100.00 | 100.00 | 100.00 | 100.00 | 100.00 | 100.00 | 100.00 | 100.00 | 100.00 | 100.00 | 100.00 |
| Te Miro Fragment | 0.00 | 0.00 | 12.67 | 2.64 | 2.64 | 100.00 | 100.00 | 0.00 | 0.00 | 0.00 | 0.00 | 100.00 | 0.00 | 0.00 | 0.00 | 100.00 | 100.00 | 0.00 | 0.00 | 0.00 |
| Te Miro North | 0.00 | 0.00 | 16.07 | 0.00 | 0.00 | 95.16 | 0.00 | 0.00 | 95.16 | 95.16 | 95.16 | 0.00 | 0.00 | 95.16 | 95.16 | 0.00 | 0.00 | 0.00 | 0.00 | 0.00 |
| Te Miro South | 0.00 | 0.00 | 16.07 | 0.00 | 0.00 | 95.16 | 0.00 | 0.00 | 95.16 | 95.16 | 95.16 | 0.00 | 0.00 | 95.16 | 95.16 | 0.00 | 0.00 | 0.00 | 0.00 | 0.00 |
| Te Tapui East | 1.37 | 0.00 | 14.54 | 4.03 | 4.03 | 0.00 | 0.00 | 0.00 | 0.00 | 0.00 | 0.00 | 0.00 | 0.00 | 0.00 | 0.00 | 0.00 | 0.00 | 0.00 | 0.00 | 0.00 |
| Te Tapui West | 1.37 | 0.00 | 14.54 | 4.03 | 4.03 | 0.00 | 0.00 | 0.00 | 0.00 | 0.00 | 0.00 | 0.00 | 0.00 | 0.00 | 0.00 | 0.00 | 0.00 | 0.00 | 0.00 | 0.00 |
| Waitomo North | 0.00 | 0.00 | 3.71 | 1.09 | 3.71 | 94.89 | 94.89 | 0.00 | 0.00 | 0.00 | 94.89 | 94.89 | 0.00 | 0.00 | 0.00 | 94.89 | 94.89 | 0.00 | 0.00 | 0.00 |
| Waitomo South | 0.00 | 0.00 | 3.71 | 1.09 | 3.71 | 94.89 | 94.89 | 0.00 | 0.00 | 0.00 | 94.89 | 94.89 | 0.00 | 0.00 | 0.00 | 94.89 | 94.89 | 0.00 | 0.00 | 0.00 |
| Walter Scott Reserve | 0.00 | 0.00 | 0.00 | 0.00 | 0.04 | 100.00 | 100.00 | 100.00 | 100.00 | 0.00 | 100.00 | 100.00 | 100.00 | 100.00 | 0.00 | 100.00 | 100.00 | 0.00 | 100.00 | 0.00 |

**Appendix S3.** (continued)

| Site | Rat controlled proportion (%) | | | | | Possum controlled proportion (%) | | | | | Rat and possum controlled proportion (%) | | | | |
| --- | --- | --- | --- | --- | --- | --- | --- | --- | --- | --- | --- | --- | --- | --- | --- |
|  | y1 | y2 | y3 | y4 | y5 | y1 | y2 | y3 | y4 | y5 | y1 | y2 | y3 | y4 | y5 |
| Hapuakohe | 0.00 | 0.00 | 0.00 | 0.00 | 0.00 | 0.00 | 0.00 | 0.00 | 0.00 | 0.00 | 0.00 | 0.00 | 0.00 | 0.00 | 0.00 |
| Kaimai North | 1.62 | 1.62 | 1.62 | 1.11 | 1.11 | 1.62 | 1.62 | 1.62 | 1.11 | 1.11 | 1.62 | 1.62 | 1.62 | 1.11 | 1.11 |
| Kaimai South | 1.62 | 1.62 | 1.62 | 1.11 | 1.11 | 1.62 | 1.62 | 1.62 | 1.11 | 1.11 | 1.62 | 1.62 | 1.62 | 1.11 | 1.11 |
| Mangatea | 100.00 | 100.00 | 0.00 | 0.00 | 0.00 | 100.00 | 100.00 | 0.00 | 0.00 | 0.00 | 100.00 | 100.00 | 0.00 | 0.00 | 0.00 |
| Maungakawa | 0.00 | 0.00 | 0.00 | 0.00 | 0.00 | 0.00 | 0.00 | 0.00 | 0.00 | 0.00 | 0.00 | 0.00 | 0.00 | 0.00 | 0.00 |
| Maungatautari Fragment | 0.00 | 0.00 | 100.00 | 100.00 | 100.00 | 0.00 | 0.00 | 100.00 | 0.00 | 0.00 | 0.00 | 0.00 | 100.00 | 0.00 | 0.00 |
| Maungatautari North 1 | 100.00 | 100.00 | 100.00 | 100.00 | 100.00 | 100.00 | 100.00 | 100.00 | 100.00 | 100.00 | 100.00 | 100.00 | 100.00 | 100.00 | 100.00 |
| Maungatautari North 2 | 100.00 | 100.00 | 100.00 | 100.00 | 100.00 | 100.00 | 100.00 | 100.00 | 100.00 | 100.00 | 100.00 | 100.00 | 100.00 | 100.00 | 100.00 |
| Owen Lewis Reserve | 0.00 | 0.00 | 0.00 | 0.00 | 0.00 | 0.00 | 0.00 | 0.00 | 39.04 | 0.00 | 0.00 | 0.00 | 0.00 | 0.00 | 0.00 |
| Pirongia East | 6.22 | 6.22 | 6.22 | 6.22 | 94.09 | 6.22 | 6.22 | 6.22 | 6.22 | 100.00 | 6.22 | 6.22 | 6.22 | 6.22 | 94.09 |
| Pirongia North | 6.22 | 6.22 | 6.22 | 6.22 | 94.09 | 6.22 | 6.22 | 6.22 | 6.22 | 100.00 | 6.22 | 6.22 | 6.22 | 6.22 | 94.09 |
| Pukemokemoke | 100.00 | 100.00 | 100.00 | 100.00 | 100.00 | 100.00 | 100.00 | 100.00 | 100.00 | 100.00 | 100.00 | 100.00 | 100.00 | 100.00 | 100.00 |
| Pureora East 1 | 18.76 | 15.36 | 10.23 | 10.29 | 0.06 | 0.00 | 0.00 | 19.70 | 0.00 | 94.94 | 0.00 | 0.00 | 0.00 | 10.29 | 0.00 |
| Pureora East 2 | 18.76 | 15.36 | 10.23 | 10.29 | 0.06 | 0.00 | 0.00 | 19.70 | 0.00 | 94.94 | 0.00 | 0.00 | 0.00 | 10.29 | 0.00 |
| Pureora South | 18.76 | 15.36 | 10.23 | 10.29 | 0.06 | 0.00 | 0.00 | 19.70 | 0.00 | 94.94 | 0.00 | 0.00 | 0.00 | 10.29 | 0.00 |
| RHMR 1 | 0.00 | 0.00 | 0.00 | 0.00 | 0.00 | 0.00 | 1.28 | 0.00 | 72.07 | 0.00 | 0.00 | 0.00 | 0.00 | 0.00 | 0.00 |
| RHMR 2 | 0.00 | 0.00 | 0.00 | 0.00 | 0.00 | 0.00 | 1.28 | 0.00 | 72.07 | 0.00 | 0.00 | 0.00 | 0.00 | 0.00 | 0.00 |
| Sanctuary Mountain | 100.00 | 100.00 | 100.00 | 100.00 | 100.00 | 100.00 | 100.00 | 100.00 | 100.00 | 100.00 | 100.00 | 100.00 | 100.00 | 100.00 | 100.00 |
| Te Miro Fragment | 100.00 | 0.00 | 0.00 | 0.00 | 0.00 | 0.00 | 100.00 | 0.00 | 0.00 | 0.00 | 0.00 | 0.00 | 0.00 | 0.00 | 0.00 |
| Te Miro North | 95.16 | 0.00 | 0.00 | 95.16 | 95.16 | 0.00 | 0.00 | 0.00 | 95.16 | 95.16 | 0.00 | 0.00 | 0.00 | 95.16 | 95.16 |
| Te Miro South | 95.16 | 0.00 | 0.00 | 95.16 | 95.16 | 0.00 | 0.00 | 0.00 | 95.16 | 95.16 | 0.00 | 0.00 | 0.00 | 95.16 | 95.16 |
| Te Tapui East | 0.00 | 0.00 | 0.00 | 0.00 | 0.00 | 0.00 | 0.00 | 0.00 | 0.00 | 0.00 | 0.00 | 0.00 | 0.00 | 0.00 | 0.00 |
| Te Tapui West | 0.00 | 0.00 | 0.00 | 0.00 | 0.00 | 0.00 | 0.00 | 0.00 | 0.00 | 0.00 | 0.00 | 0.00 | 0.00 | 0.00 | 0.00 |
| Waitomo North | 0.00 | 0.00 | 0.00 | 0.00 | 0.00 | 94.89 | 94.89 | 0.00 | 0.00 | 0.00 | 0.00 | 0.00 | 0.00 | 0.00 | 0.00 |
| Waitomo South | 0.00 | 0.00 | 0.00 | 0.00 | 0.00 | 94.89 | 94.89 | 0.00 | 0.00 | 0.00 | 0.00 | 0.00 | 0.00 | 0.00 | 0.00 |
| Walter Scott Reserve | 100.00 | 100.00 | 100.00 | 0.00 | 0.00 | 100.00 | 100.00 | 100.00 | 100.00 | 0.00 | 100.00 | 100.00 | 100.00 | 0.00 | 0.00 |

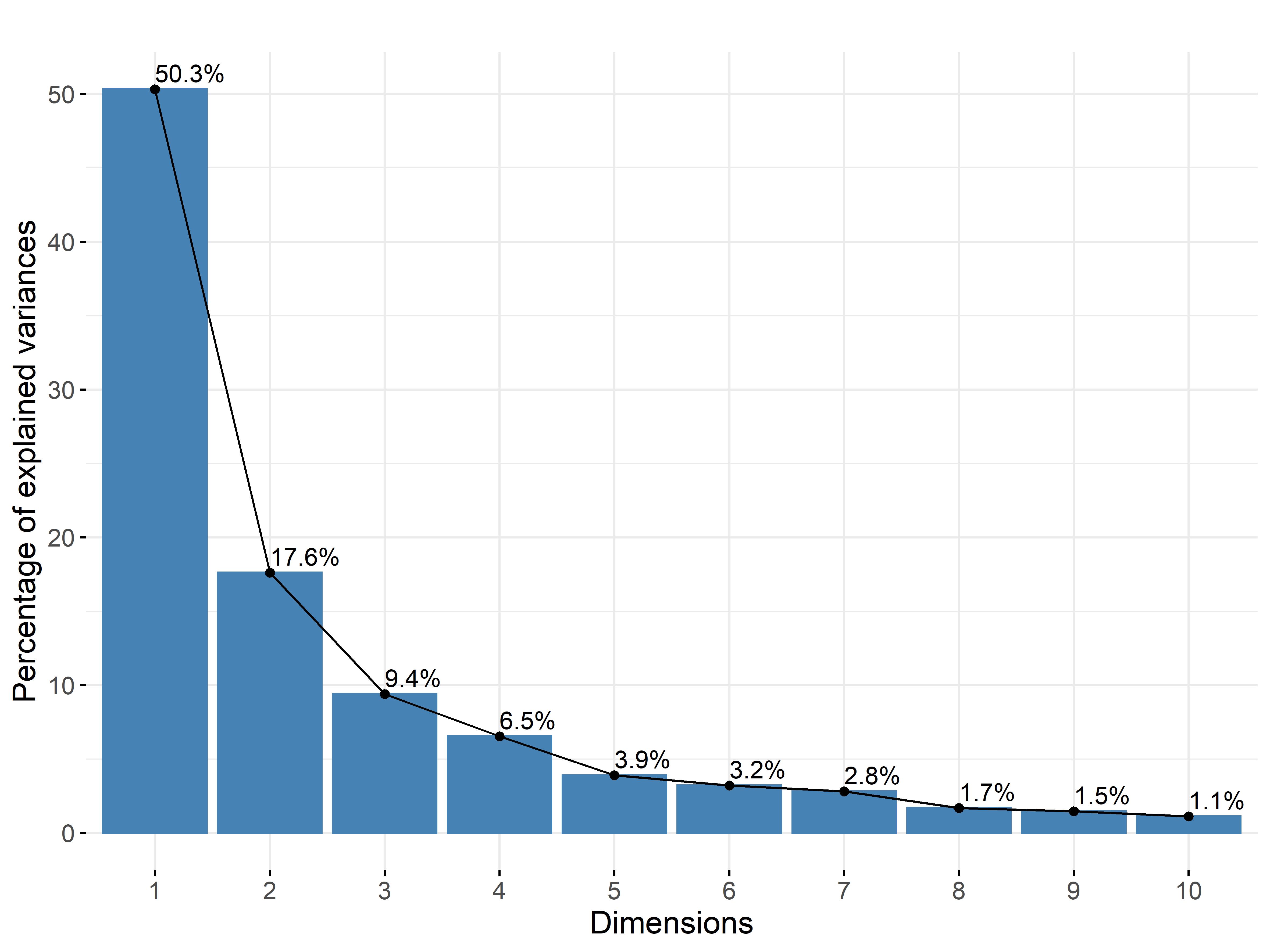

**Figure S4.** Scree plot showing the proportion of variance within the invasive predator control dataset (Appendix 3) explained by each principle component (dimension) generated through principle components analysis.

**Appendix S5.** Traits and abundances of native New Zealand birds recorded in this study. Weight, clutch size, lifespan, breeding age, feeding guild and nesting habitat data were obtained from New Zealand Birds Online (Miskelly, n.d.). Gap-crossing ability describes the potential ability of each species to move across pasture* and water** gaps between forest habitat based on the estimates of Burge, Innes, Fitzgerald, & Richardson (2017). As no estimates of the gap-crossing ability of Grey Warblers exists, natal dispersal*** distance was used as a rough approximation. Functional groups were determined from a Gower dissimilarity matrix based on trait and site-specific abundance data (Appendix 6).

| Species | Weight (g) | Gap-crossing ability (km) | Clutch size | Lifespan (years) | Breeding age (months) | Primary feeding guild | Nesting habitat | Abundance | Functional group |
| --- | --- | --- | --- | --- | --- | --- | --- | --- | --- |
| New Zealand Bellbird (*Anthornis melanura*) | 30 | 22** | 4 | 8 | 12 | Nectarivore | Arboreal | 10 | 1 |
| New Zealand Fantail (*Rhipidura fuliginosa*) | 8 | 0.15** | 3.5 | 5 | 12 | Insectivore | Arboreal | 184 | 2 |
| Grey Warbler (*Gerygone igata*) | 6.5 | 0.9*** | 3.5 | 5 | 12 | Insectivore | Arboreal | 54 | 2 |
| Rifleman (*Acanthisitta chloris*) | 6.5 | 0.3* | 3.5 | 9 | 6 | Insectivore | Arboreal | 15 | 2 |
| North Island Robin (*Petroica longipes*) | 35 | 0.11* | 2.45 | 3 | 12 | Insectivore | Arboreal | 25 | 2 |
| New Zealand Tomtit (*Petroica macrocephala*) | 11 | 0.12** | 3.9 | 3 | - | Insectivore | Arboreal | 20 | 2 |
| Tui (*Prosthemadera novaeseelandiae*) | 105 | 20* | 3 | 12 | 24 | Nectarivore | Arboreal | 101 | 1 |
| Waxeye (*Zosterops lateralis*) | 13 | 0.16** | 2.5 | 9 | 10 | Omnivore | Arboreal | 15 | 3 |
| New Zealand wood pigeon (*Hemiphaga novaeseelandiae*) | 650 | 33** | 1 | 21 | - | Herbivore | Arboreal | 14 | 4 |

**Appendix S6.** Native bird abundances at the 26 study sites. Abundance values are the sum of three 5-minute bird counts performed at each site from April to July of 2019.

| Site | New Zealand Bellbird (Anthornis melanura) | New Zealand Fantail (Rhipidura fuliginosa) | Grey Warbler (Gerygone igata) | Rifleman (Acanthisitta chloris) | North Island Robin (Petroica longipes) | New Zealand Tomtit (Petroica macrocephala) | Tui (Prosthemadera novaeseelandiae) | Waxeye (Zosterops lateralis) | New Zealand wood pigeon (Hemiphaga novaeseelandiae) |
| --- | --- | --- | --- | --- | --- | --- | --- | --- | --- |
| Hapuakohe | 0 | 6 | 1 | 0 | 0 | 0 | 0 | 1 | 0 |
| Kaimai North | 0 | 13 | 4 | 0 | 0 | 0 | 1 | 0 | 0 |
| Kaimai South | 1 | 12 | 2 | 0 | 0 | 0 | 5 | 0 | 0 |
| Mangatea | 0 | 9 | 0 | 0 | 0 | 1 | 0 | 0 | 1 |
| Maungakawa | 0 | 5 | 4 | 0 | 0 | 0 | 4 | 0 | 0 |
| Maungatautari Fragment | 0 | 9 | 4 | 0 | 0 | 0 | 6 | 0 | 1 |
| Maungatautari North 1 | 1 | 5 | 0 | 0 | 7 | 6 | 3 | 0 | 0 |
| Maungatautari North 2 | 3 | 7 | 2 | 0 | 3 | 5 | 10 | 0 | 2 |
| Owen Lewis Reserve | 0 | 10 | 1 | 0 | 0 | 0 | 3 | 0 | 0 |
| Pirongia East | 0 | 7 | 2 | 0 | 0 | 0 | 11 | 0 | 1 |
| Pirongia North | 0 | 4 | 2 | 2 | 0 | 0 | 3 | 0 | 1 |
| Pukemokemoke | 0 | 11 | 1 | 0 | 0 | 0 | 4 | 0 | 0 |
| Pureora East 1 | 0 | 3 | 1 | 0 | 0 | 0 | 2 | 1 | 1 |
| Pureora East 2 | 1 | 12 | 1 | 1 | 2 | 0 | 3 | 3 | 0 |
| Pureora South | 0 | 4 | 1 | 1 | 4 | 1 | 3 | 0 | 3 |
| RHMR 1 | 0 | 2 | 0 | 0 | 1 | 1 | 2 | 0 | 0 |
| RHMR 2 | 1 | 3 | 0 | 1 | 0 | 1 | 0 | 1 | 0 |
| Sanctuary Mountain | 3 | 10 | 1 | 2 | 8 | 4 | 7 | 0 | 0 |
| Te Miro Fragment | 0 | 7 | 5 | 0 | 0 | 0 | 3 | 6 | 1 |
| Te Miro North | 0 | 4 | 6 | 0 | 0 | 0 | 1 | 0 | 1 |
| Te Miro South | 0 | 13 | 2 | 2 | 0 | 0 | 4 | 1 | 1 |
| Te Tapui East | 0 | 8 | 2 | 0 | 0 | 0 | 10 | 0 | 1 |
| Te Tapui West | 0 | 3 | 1 | 0 | 0 | 0 | 4 | 0 | 0 |
| Waitomo North | 0 | 8 | 2 | 3 | 0 | 0 | 5 | 2 | 0 |
| Waitomo South | 0 | 4 | 4 | 2 | 0 | 0 | 2 | 0 | 0 |
| Walter Scott Reserve | 0 | 5 | 5 | 1 | 0 | 1 | 5 | 0 | 0 |

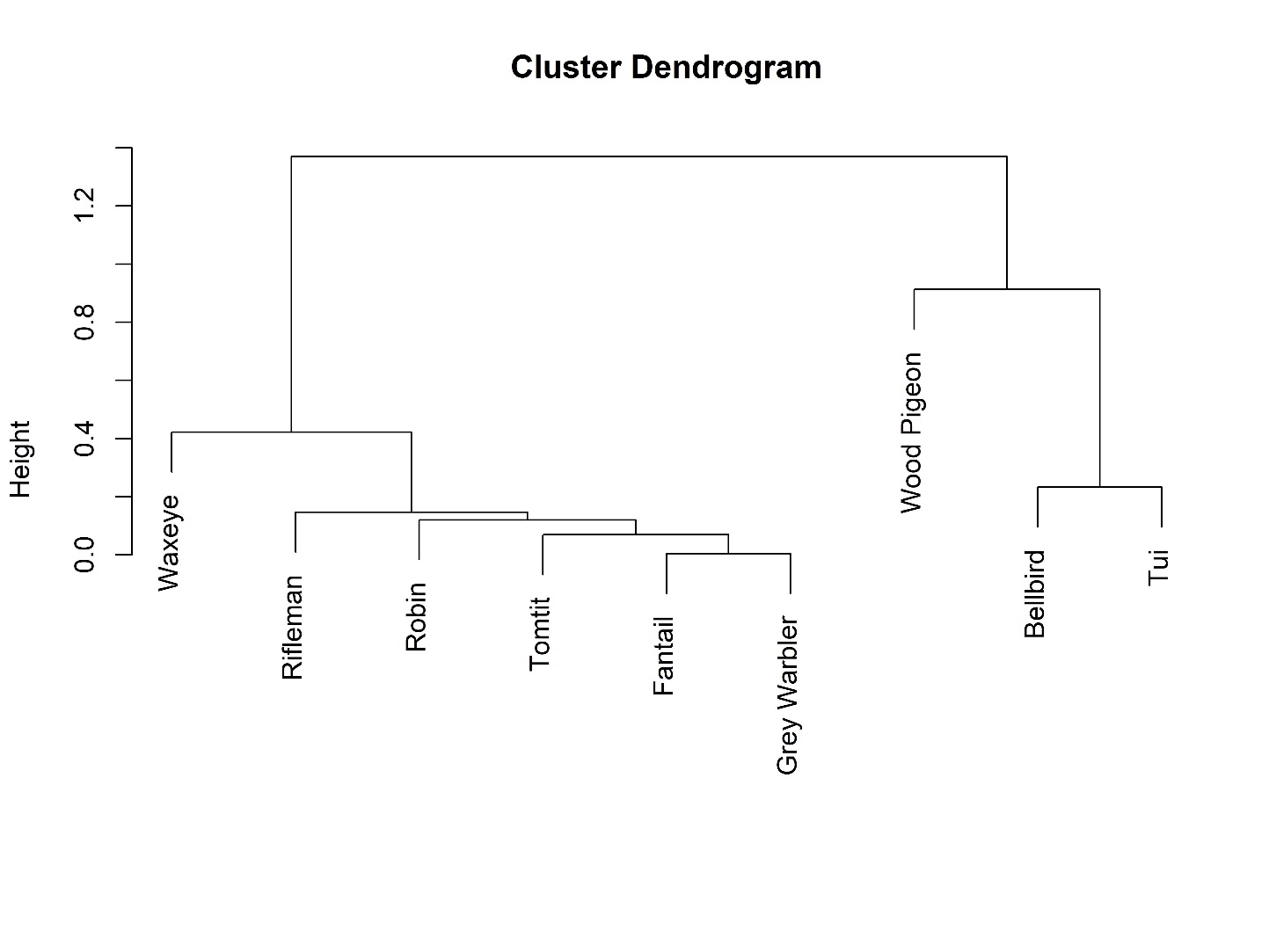
**Appendix S7.** Dendrogram showing the hierarchical relationships between native bird species based on a Gower dissimilarity matrix created from species specific trait data (Appendix S5) weighted by abundance counts (Appendix S6).

**Appendix S8.** Statistics for generalised linear models with negative binomial distributions containing individual invasive predator control predictor variables.

| Response variable | Predictor variable | Intercept | Slope | z-score | p-value |
| --- | --- | --- | --- | --- | --- |
| Total invasive predator abundance | Control intensity | -0.899 | -0.935 | -4.545 | < .001 |
| Total invasive predator abundance | Temporal distribution of control | -0.660 | 0.011 | 0.057 | 0.954 |
| Brushtail possum abundance | Control intensity | -1.865 | -1.013 | -3.021 | 0.003 |
| Brushtail possum abundance | Temporal distribution of control | -1.910 | 0.302 | 0.991 | 0.322 |
| Ship rat abundance | Control intensity | -2.119 | -0.672 | -1.927 | 0.054 |
| Ship rat abundance | Temporal distribution of control | -2.084 | -0.337 | -1.157 | 0.247 |

**Appendix S9.** Confidence intervals (95%) for predictor variables in averaged models for invasive predator and native bird abundance models. Intervals which do not included zero are indicated in bold text.

| Predictor | Total invasive predator abundance | Brushtail possum abundance | Ship rat abundance | Total native bird abundance | Nectarivore abundance | Insectivore abundance |
| --- | --- | --- | --- | --- | --- | --- |
| Control intensity | **-1.421 – -0.509** | **-1.652 – -0.461** | -0.705 – 0.454 | **0.066 – 0.359** | -0.147 – 0.197 | **0.151 – 0.412** |
| Temporal distribution of control | -0.075 – 0.077 | -0.294 – 0.691 | -0.188 – 0.159 | -0.041 – 0.041 | -0.237 – 0.145 | -0.039 – 0.040 |
| Area | -0.294 – 0.210 | -0.228 – 0.170 | -0.118 – 0.108 | **0.009 – 0.307** | -0.155 – 0.252 | **0.051 – 0.381** |
| Fractal dimension index | -0.124 – 0.124 | -0.314 – 0.232 | **0.174 – 1.696** | -0.267 – 0.113 | **-0.744 – -0.014** | -0.122 – 0.090 |
| Exotic forest % (1-5 km) | -0.124 – 0.107 | -0.196 – 0.173 | -0.584 – 0.334 | -0.049 – 0.052 | -0.119 – 0.151 | -0.057 – 0.050 |
| Native forest % (1-5 km) | -0.091 – 0.092 | -0.136 – 0.133 | -1.831 – 0.216 | -0.105 – 0.081 | -0.199 – 0.150 | -0.045 – 0.045 |
| Exotic forest % (< 1 km) | -0.179 – 0.235 | -0.151 – 0.169 | -0.308 – 0.214 | -0.201 – 0.090 | -0.130 – 0.194 | -0.316 – 0.056 |
| Native forest % (< 1 km) | -0.093 – 0.101 | -0.222 – 0.180 | -0.139 – 1.850 | -0.154 – 0.087 | -0.192 – 0.139 | -0.070 – 0.060 |
